# Test-retest reliability of MEG functional brain connectivity related to language processing

**DOI:** 10.1101/2023.09.27.559879

**Authors:** Heidi Ala-Salomäki, Marijn van Vliet, Jan Kujala, Timo Roine, Mia Liljeström, Riitta Salmelin

**Affiliations:** Department of Neuroscience and Biomedical Engineering, Aalto University, FI-00076 Aalto, Finland; Aalto NeuroImaging, Aalto University, FI-00076 Aalto, Finland; Department of Psychology, University of Jyväskylä, FI-40014, Finland; BioMag Laboratory, HUS Medical Imaging Center, Helsinki University Hospital, Helsinki University and Aalto University School of Science, Helsinki, FI-00290, Finland

**Keywords:** functional connectivity, structural connectivity, magnetoencephalography, picture naming

## Abstract

The number of studies examining changes in functional connectivity of the human brain is increasing rapidly. In this magnetoencephalography (MEG) study, we examined the reliability of dynamic connectivity related to language processing in a picture naming test-retest paradigm, using data collected from the same participants on two separate days. We determined the connections that were reliable across both days and also examined the behavioral, functional, and structural properties underlying this reliability. A particularly salient finding among a rich set of results was a reliable pattern of beta connectivity increase in the left motor and frontal regions (0–400 ms and 400–800 ms after stimulus onset) and gamma connectivity decrease in the bilateral motor regions (800–1200 ms) which we suggest to represent the motor preparation of speech production. Furthermore, the reliable connections tended to be more frequently associated with the behavioral performance than the non-reliable ones. Finally, the reliable connections were also linked to stronger functional connectivity, as well as to stronger structural connectivity and shorter structural path length, as determined through diffusion MRI (magnetic resonance imaging). Overall, this study defines reliable language-related functional connectivity and introduces practices that may increase reliability.

**AUTHOR SUMMARY:** Research applying connectivity metrics in neuroimaging has increased rapidly during recent years. Hence, the focus has also been to define the best methods for increasing the reliability of connectivity estimation. This study determined reliable functional connectivity from MEG data related to language processing. Moreover, we defined what makes a connection reliable by studying the behavioral, functional, and structural properties underlying the reliable connections.

## 1 INTRODUCTION

The use of functional connectivity metrics in neuroscience research is rapidly growing, which has also highlighted the importance of the reliability and validity of those metrics (see e.g., Noble, Scheinost and Constable, 2019). Reliability refers to the consistency of a measure across repeated measurement sessions, and validity refers to the association of a measure with a defined process. In this study, we sought to determine the reliability of magnetoencephalography (MEG) functional connections in a test-retest scenario involving a language task.

Many studies of functional connectivity are performed on resting-state data. However, rest may not be the optimal condition for obtaining reliable connectivity estimates, since its unconstrained nature can lead to non-stationary signal patterns due to e.g., arousal and movement (Cohen 2018; Elton and Gao, 2015; Finn et al., 2017; Greene, Gao, Scheinost and Constable, 2018; Noble et al., 2019), which may cause spurious connections that do not replicate in a test-retest scenario (reviewed in Finn et al., 2017; Noble et al., 2019). Furthermore, resting-state connectivity analysis typically provides a static view of the mean connectivity patterns over time. Yet, functional connectivity is thought to reconfigure in meaningful ways across time and task states (Cohen, 2018; Elton and Gao, 2015; Medaglia, Lynall and Bassett, 2015). An event-related study design that evokes repeatable task states may therefore be beneficial for reliability. Hence, we examined the test-retest reliability during a picture naming task, which is a constrained, yet engaging language task that requires the interaction between multiple neural systems. By contrasting the picture naming task with a simple visual task, we aimed to highlight language-related functional networks.

For language tasks, earlier studies (Chai, Mattar, Blank, Fedorenko and Bassett, 2016; DeSalvo, Douw, Takaya, Liu and Stufflebeam, 2014; Li et al., 2020; Liljeström, Kujala, Stevenson and Salmelin, 2015; Liljeström, Vartiainen, Kujala and Salmelin, 2018; Schoffelen, Hultén, Marquand, Uddén and Hagoort, 2017; Tomasi and Volkow, 2020) have identified language-specific connectivity patterns. In earlier language studies, the focus has typically been on a limited selection of activation derived or user-defined brain regions or restricted exclusively to the left hemisphere (see e.g. Schoffelen et al., 2017). However, the core functional connections for language processing remain unknown and may reach outside of areas typically associated with increased activity. For a comprehensive description of the connectivity patterns related to language, we chose to widen the spatial perspective to include both hemispheres and also brain regions that are not commonly reported in language studies focusing on local changes of cortical activity.

In addition to determining the reliability of the connectivity estimation, the validity of the observed connectivity patterns needs to be verified. One approach to validate the functional connectivity patterns is to relate them to behavior, since connections linked to behavior are likely to represent task-relevant brain processes (Cohen, 2018; Jiang et al., 2020). Therefore, an increasing number of studies have begun to investigate the relationship of functional connectivity to behavioral performance, also regarding language function (Mollo et al., 2016; Tomasi and Volkow, 2020; Vatansever et al., 2017; Yu et al., 2018).

There are also other factors that might influence the reliability of the functional connections. Earlier findings suggest that the strength of a functional connection predicts its reliability (Garcés, Martín-Buro and Maestú, 2016; Noble et al., 2019). There is also some evidence that structurally connected brain regions have more reliable functional connections than structurally unconnected ones (Griffa et al., 2017; Shen, Hutchison, Bezgin, Everling and McIntosh, 2015). Accordingly, beyond determining whether functional connections were reliable, we investigated how the observed reliability linked to the subjects’ behavior and the strength of both functional and structural connectivity.

The present MEG study aims to determine reliable dynamic functional connections related to language processing in a picture naming test-retest paradigm. Within the picture naming task, we focused on the processing stages prior to overt speech production. We combined a well-controlled study design with a data-driven approach that did not establish anatomical constraints *a priori*. The field of neuroimaging has begun to define best methods for increasing reliability (see for a review Noble et al., 2019), and this study expands upon them by studying the behavioral, functional, and structural properties behind the reliable connections.

## 2 METHODS

### 2.1 Participants

Twenty healthy human participants took part in the MEG measurements (10 females and 10 males; mean age 25 years; SD 3.9; age range 21–35 years). MEG data was recorded in two separate measurement sessions (1–13 days apart, mean 4.2, SD 3.9). All participants were native Finnish speakers, right-handed, had no history of neurological, psychiatric, or developmental disorders or learning disabilities, and had normal or corrected-to-normal vision. One participant’s data was not used for the analysis due to non-compliance with the MEG task instruction. In addition, one participant was excluded from the analysis of the behavioral data due to non-compliance with the behavioral task and another participant from the analysis of structural data due to preprocessing difficulties. A written informed consent was obtained from all participants, in agreement with the prior approval of the Aalto University Research Ethics Committee. The MEG evoked responses and oscillatory activity of this data set have previously been analysed in Ala-Salomäki, Kujala, Liljeström and Salmelin (2021).

### 2.2 Experimental design

#### 2.2.1 Tasks

In the naming task, pictures were presented to the participants, and the participants overtly named the object in the picture. If the participant did not know the name of the object, they were instructed to say “skip” (“ohi” in Finnish). In the visual task, participants said “yes” (“kyllä” in Finnish) when a target picture was presented (a red cross in the middle, on top of the picture). The participants also performed other tasks that were not analysed in the present study (see Ala-Salomäki et al. 2021). Outside the MEG device the participants performed the following behavioral tasks: The Boston naming task (BNT), Rapid automatized naming (RAN), Rapid alternating stimulus (RAS), Fluency animal, and Fluency s-letter.

#### 2.2.2 Stimuli and procedures

The naming task and the visual task each included 100 different pictures of objects. Half of the pictures (set A) were presented in one measurement session and the other half (set B) in another measurement session. Each picture was presented twice during the measurement session. Half of the participants started with set A and the other half with set B. The properties of the pictures are described in detail in Ala-Salomäki et al. (2021). The visual task, in each measurement session, also included 20 target stimuli (not included in the analysis). The picture presentation protocol was the same in both tasks and measurement sessions. Pictures were presented to the participants in blocks, and only one task was performed during one block. Each block contained ten pictures. For the visual task, each block included 1–3 target stimuli. In the block, first a fixation cross appeared on the screen for 1 s, followed by the picture for 300 ms. Then, a blank screen was presented for 1 s, followed by a question mark for 2 s. Participants gave their answer, when the question mark was presented. A measurement session included 10 blocks for the naming and 12 blocks for the visual task. The order of the blocks was randomized.

### 2.3 MEG and (diffusion) MRI recordings

MEG data was measured at the Aalto NeuroImaging MEG Core (Aalto University, Espoo, Finland) using a Vectorview whole-head MEG device (MEGIN, Helsinki, Finland). The device comprises 306 sensors: 204 planar gradiometers and 102 magnetometers, each coupled to a Superconducting Quantum Interference Device (SQUID) in a helmet-shaped array. Electro-oculogram signals were recorded for identifying eye blinks and saccades. The MEG signals were filtered at 0.1–330 Hz and sampled at 1000 Hz. Details on the MEG recordings are described in Ala-Salomäki et al. (2021). MRIs were acquired at the Aalto NeuroImaging Advanced Magnetic Imaging Centre using a Siemens Magnetom Skyra 3.0 T MRI scanner. T1-weighted images were acquired with a standard gradient echo sequence and 1 mm ✕ 1 mm ✕ 1 mm resolution. Diffusion MRI data were acquired with a voxel size of 2 mm ✕ 2 mm ✕ 2 mm in 64 uniformly distributed gradient orientations and a diffusion weighting of b=1000 s/mm2. A total of five b=0 s/mm2 images were acquired at the beginning and end of the sequence. The full acquisition was repeated in the reverse phase-encoding direction, resulting in a total of 138 volumes.

### 2.4 MEG preprocessing and connectivity analysis

The spatiotemporal signal space separation method (Taulu and Simola, 2006) was applied to MEG data for noise reduction (16 s temporal window, a subspace correlation limit of 0.98, inside expansion order of 8, and outside expansion order of 3). The head position data was used to transform the MEG data to a common position (MEGIN Maxfilter software package, 2.2.12). Independent Component Analysis was applied to remove eye-blink artifacts. In the naming task, only epochs with correct responses were analyzed.

For estimating all-to-all MEG functional connectivity, we applied a pipeline (van Vliet, Liljeström, Aro, Salmelin and Kujala, 2018) that uses coherence as the connectivity metric, estimated utilizing the Dynamic Imaging of Coherent Sources (DICS) spatial filter originally proposed by Gross et al. (2001), combined with a wavelet approach to achieve a high temporal resolution (Kujala, Vartiainen, Laaksonen and Salmelin, 2012; Laaksonen, Kujala and Salmelin, 2008). Connectivity was analyzed for the naming and the visual task in multiple frequency bands and time windows, separately for the two measurement sessions. Data were analyzed in the following frequency bands of interest: 4–7 Hz (theta), 8–13 Hz (alpha), 14–20 Hz (low beta), 21–30 Hz (high beta), 31–45 Hz (low gamma), and 60–90 Hz (high gamma), similarly to Ala-Salomäki et al. (2021). Time windows 0–400 ms, 400–800 ms, and 800–1200 ms time-locked to stimulus presentation were selected similarly to Ala-Salomäki et al. (2021). Cross-spectral density matrices (CSDs) across planar gradiometer channels were calculated for the two tasks in the selected time windows using time-frequency representation (Morlet wavelets) in the frequency range 4–90 Hz, at 1-Hz intervals for the 4–13 Hz frequency range and 2-Hz intervals for the 14–90 Hz frequency range. CSD matrices were transformed into a cortical representation using the spatial filter.

A surface-based source space in the FreeSurfer (Dale, Fischl and Sereno, 1999) template brain (‘fsaverage’) was created, and these source locations were transformed to each individual subject. We used a regularly spaced grid of 5,124 points on each hemisphere, with an average distance of 2.6 mm between neighbouring points from which we excluded points that were further than 7 cm from the closest MEG sensor in an example subject. Coherence was computed between each anatomical grid point and all other grid points, using spherical tangential source orientations that yielded maximum coherence. Coherence was estimated separately for each measurement session, task, frequency band, and time window of interest. To avoid spurious connectivity between closely situated areas due to spatial leakage (field spread), a minimum distance limit of 4 cm between the connection start and end points was applied, and coherence values were always evaluated by contrasting two experimental conditions to cancel out spatial leakage that is constant across conditions (see also section 2.6 Power difference between the tasks). For more details, see van Vliet et al. (2018).

### 2.5 Parcellation

For further analysis of the connectivity data, a custom-made parcellation based on the Destrieux FreeSurfer template parcellation for ‘fsaverage’ was used. The custom-made parcellation was constructed to produce as similar sized parcels as possible to overcome the need of degree correction, while preserving coarse anatomical boundaries when possible. The parcellation included 55 brain regions per hemisphere (see Fig. 1). The details of the parcellation are described in Ala-Salomäki et al. (2021). For graph-theoretical analyses of diffusion MRI data, subcortical structures segmented with FIRST in FMRIB Software Library (FSL) were included (Jenkinson, Beckmann, Behrens, Woolrich and Smith, 2012).

**Figure 1.**
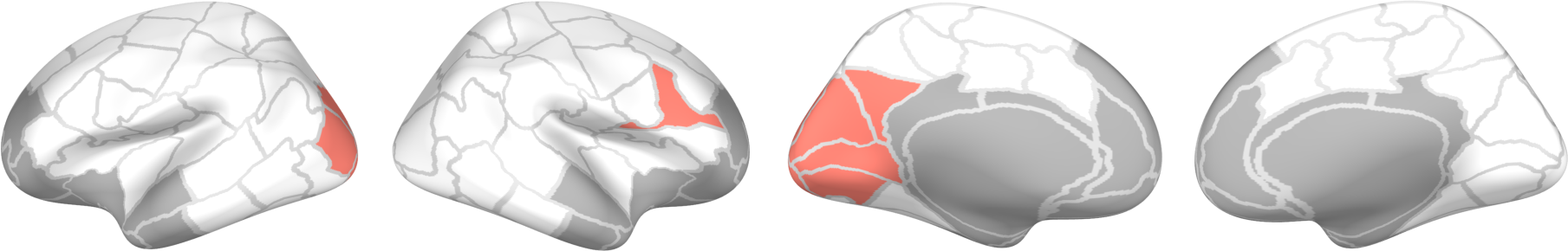
Parcellation and power differences between the tasks. Parcels that were used in the analysis. Gray parcels represent the anterior and subcortical parcels that were excluded from the analysis. Red parcels represent the parcels, where the power difference between the naming and the visual task was larger than 20 % for more than four subjects in individual time-frequency bands and measurement sessions.

### 2.6 Power difference between the tasks

Power differences between the experimental tasks can introduce spurious differences in the coherence estimates (Gross et al., 2013; Kujala et al., 2007; Kujala, Gross, and Salmelin, 2008). Therefore, power levels were computed from the CSD matrices described above for the tasks, frequency bands, and time windows of interest, separately for the two measurement sessions. The power differences were further computed between the naming and the visual task at the parcel level for each subject. We determined the parcels where the power difference between the tasks was larger than 20 % for more than four subjects (Figure 1). Power differences were detected in only a few parcels (mainly non-overlapping frequency bands and time windows) and were not present in both measurement sessions. Therefore, we considered that the data did not show potentially confounding power differences, and no frequency band was excluded from the further analysis.

### 2.7 Diffusion MRI preprocessing and connectivity analysis

Diffusion MRI data were corrected for subject motion (Leemans and Jones, 2009), eddy current induced distortions (Andersson and Sotiropoulos, 2016) and susceptibility induced distortions by using FSL (Jenkinson et al., 2012). Correction for B1 field inhomogeneities was performed using N4BiasFieldCorrection in Advanced Normalization Tools (Avants, Tustison and Song, 2009; Tustison et al., 2010) and MRtrix3 (Tournier et al., 2019). Statistical parametric mapping was used to rigidly coregister the T1-weighted images to the corrected diffusion MRI data (Collignon et al., 1995; Penny, Friston, Ashburner, Kiebel and Nichols, 2011).

We used Constrained Spherical Deconvolution to estimate voxel-level fiber orientation distributions with maximum 8th order spherical harmonics in MRtrix3 (Tournier, Calamante and Connelly, 2007; Tournier et al., 2019). We performed whole-brain probabilistic streamlines tractography with the iFOD2 algorithm (Tournier et al., 2009) by seeding from the gray matter–white matter interface and using anatomically-constrained tractography to ensure anatomical validity of the streamlines (Smith, Tournier, Calamante and Connelly, 2012). Spherical-deconvolution informed filtering of tractograms was used to correct for the density bias in the reconstructed whole-brain streamlines by iteratively optimizing the streamline weights to match the local fiber orientation distributions (Smith, Tournier, Calamante and Connelly, 2013; Smith, Tournier, Calamante and Connelly, 2015). Connectivity matrices were weighted by the sum of streamline weights connecting a pair of gray matter regions. Previously, the test-retest reliability of the structural brain connectivity networks has been shown to be high (Roine et al., 2019). The shortest path length between a pair of nodes and edge betweenness centrality were calculated using the Brain Connectivity Toolbox (Rubinov and Sporns, 2010).

### 2.8 Consistency analysis of MEG connectivity

#### 2.8.1 Statistical evaluation of naming-related connectivity for selecting parcel-level connections to be included in the consistency analysis

For identifying naming-related connections, we evaluated the connectivity modulations between the naming and the visual task in the first measurement session. We conducted paired t-tests for all connections at two significance levels (p<0.001 and p<0.0001) for the selected frequency bands and time windows. We did not include the most anterior areas (see gray areas in Figure 1) in the analysis, as signals and, subsequently, connectivity of those areas could represent artifactual activity related to eye movements. We thresholded the connectivity results using clustering, such that significant connections were grouped into bundles. Bundles with a minimum of 20 connections and a maximum distance of 1.3 cm between connections were retained. Then we defined these significant naming-related connections at the parcel level. The resulting parcel-level connections (all possible connections within parcels) for each p-value, frequency band and time window served as input for the subsequent consistency analysis. The modulation direction of connectivity (increase or decrease) was determined by taking the group mean of the connectivity difference between the naming and the visual task during the first measurement session at the parcel level (all possible connections within parcels).

#### 2.8.2 ICC

To establish naming-related connections that were consistent across measurement sessions, we performed intraclass correlation coefficient (ICC) analysis on the predefined parcel-level connections separately for each p-value, frequency band and time window (see 2.8.1 Statistical evaluation of naming-related connectivity for selecting parcel-level connections to be included in the consistency analysis). To compute the ICC value for a specific connection at the parcel level, each subject’s connectivity difference between the naming and the visual task was taken and then summed at the parcel level (all possible connections within parcels) separately for the two measurement sessions.

ICC is defined as:

ICC(3, 1) = (BMS-EMS)/(BMS+(k-1)*EMS),

Where BMS = between-subjects mean square, EMS = error mean square, and k = number of repeated sessions.

With the consistent connections we refer to connections with ICC>0.4 (Cicchetti, 1994). With the inconsistent connections we refer to connections with ICC≤0.4.

### 2.9 Properties behind the consistent connections

We studied the properties behind the consistent connections by comparing the consistent connections to the inconsistent ones. For that purpose, data across frequency bands and time windows was pooled together.

#### 2.9.1 Behavioral performance and consistency of connectivity

We investigated whether the behavioral performance explained functional connectivity strength more often for the consistent or the inconsistent connections. This was done by a multiple regression model for each connection (no p-value correction, several multiple regression analyses done). Each subject’s behavioral performances (combined mean variable of RAN and RAS, fluency s, and fluency animal) were set as independent variables. Each subject’s mean of MEG connectivity difference between the naming and the visual task of both days were set as dependent variables. A chi-square test was used to test whether the regression model fitted to the data more often in the consistent or the inconsistent connections.

#### 2.9.2 MEG connectivity strength and consistency of connectivity

We investigated whether the MEG-derived connectivity strength was systematically higher for the consistent or the inconsistent connections. For each connection, each subject’s connectivity difference between the naming and the visual task was obtained and then summed at the parcel level separately for the two measurement sessions, after which the absolute mean across participants and measurement days was calculated. A non-parametric Mann-Whitney U-test was used to test whether the MEG connectivity strength was different between the consistent and the inconsistent connections.

#### 2.9.3 Structural number of direct streamlines, shortest path length, and betweenness centrality related to consistency of functional connectivity

We investigated whether the number of direct streamlines, connection’s shortest path length, and connection’s betweenness centrality differed between the consistent and the inconsistent connections. For each MEG-derived connection, the mean number of direct streamlines, the mean of shortest path length, and the mean of betweenness centrality across participants was calculated. A non-parametric Mann-Whitney U-test was used to test whether these properties were different between the consistent and the inconsistent connections.

## 3 RESULTS

### 3.1 Effect of statistical threshold on the number and the percentage of the consistent connections

To select naming-related connections, we calculated the connectivity modulation between the naming and the visual task at significance levels of p<0.001 and p<0.0001 for each frequency band and time window. These connections served as input to the ICC analysis. When pooling together all the frequency bands and time windows, a total of 5054 (statistical threshold p<0.001) and 514 (statistical threshold p<0.0001) connections were selected. In the subsequent ICC analysis, 8.9 % (statistical threshold p<0.001) and 11.5 % (statistical threshold p<0.0001) of those connections were consistent. Figure 2 illustrates how the choice of statistical threshold influences the number of the consistent connections and the percentage of the consistent connections in a specific frequency band and time window: the percentages of the consistent connections were up to 12.7 % (statistical threshold p<0.001) and 29.2 % (statistical threshold p<0.0001).

**Figure 2.**
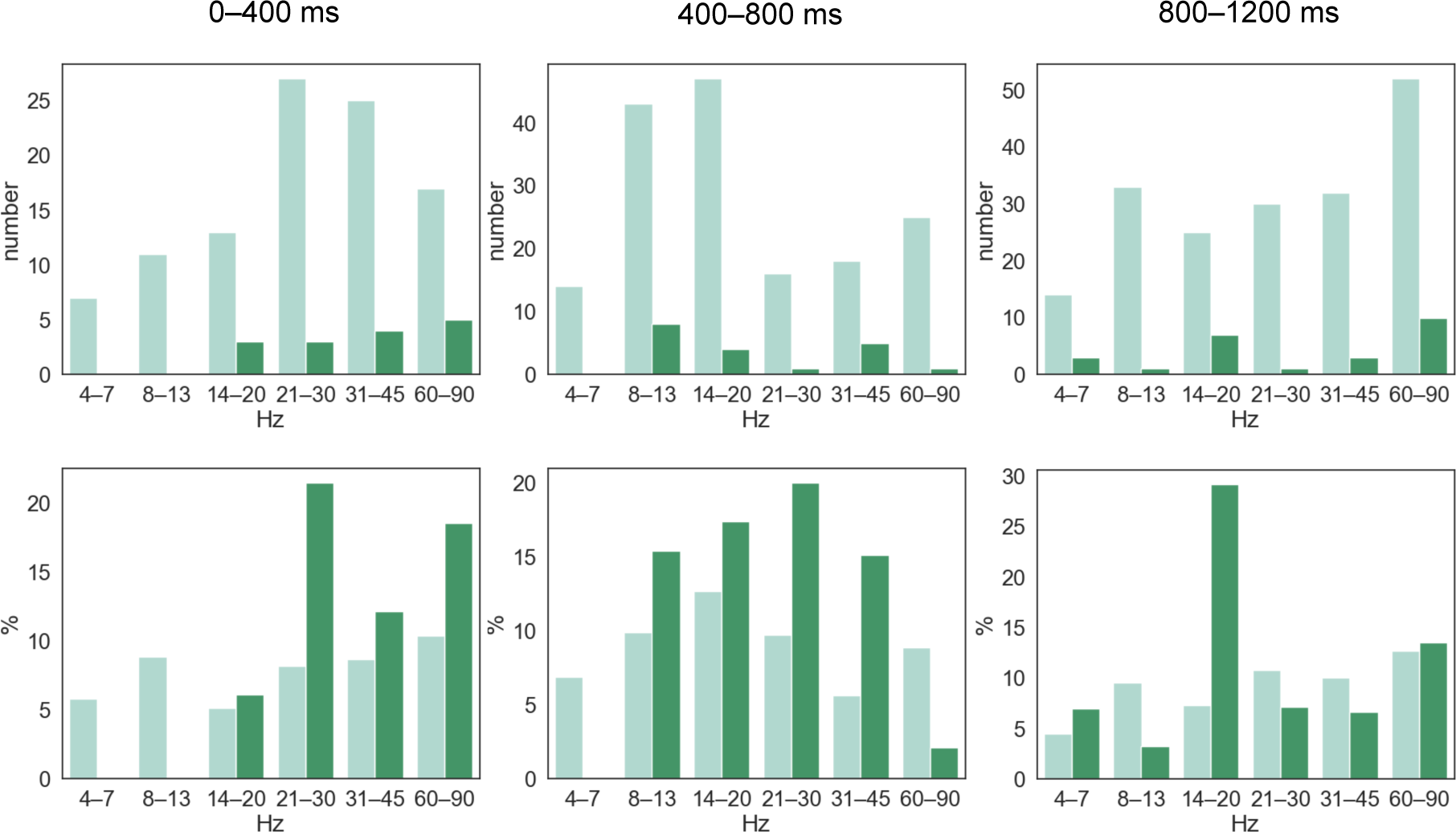
Numbers and percentages of the consistent connections. The numbers of the consistent connections and the percentages of the consistent connections of all the selected connections at different statistical thresholds (p<0.001 light green; p<0.0001 dark green), separately for each frequency band and time window. Zero means that no connection was selected for the ICC analysis.

### 3.2 Consistent connections related to picture naming

Increased connectivity in the picture naming task compared to the visual task was observed consistently and dynamically in different time windows, frequency bands and hemispheres as illustrated in Figure 3. In the early time window (0–400 ms), naming-related gamma connectivity increased consistently in the visual and posterior temporal regions. From the early (0–400 ms) to the middle (400–800 ms) time window, consistent left-lateralized naming-related beta connectivity increase was observed mainly in the motor and frontal regions. In the late (800–1200 ms) time window naming-related theta connectivity increased consistently in the motor, frontal, and insular regions.

**Figure 3.**
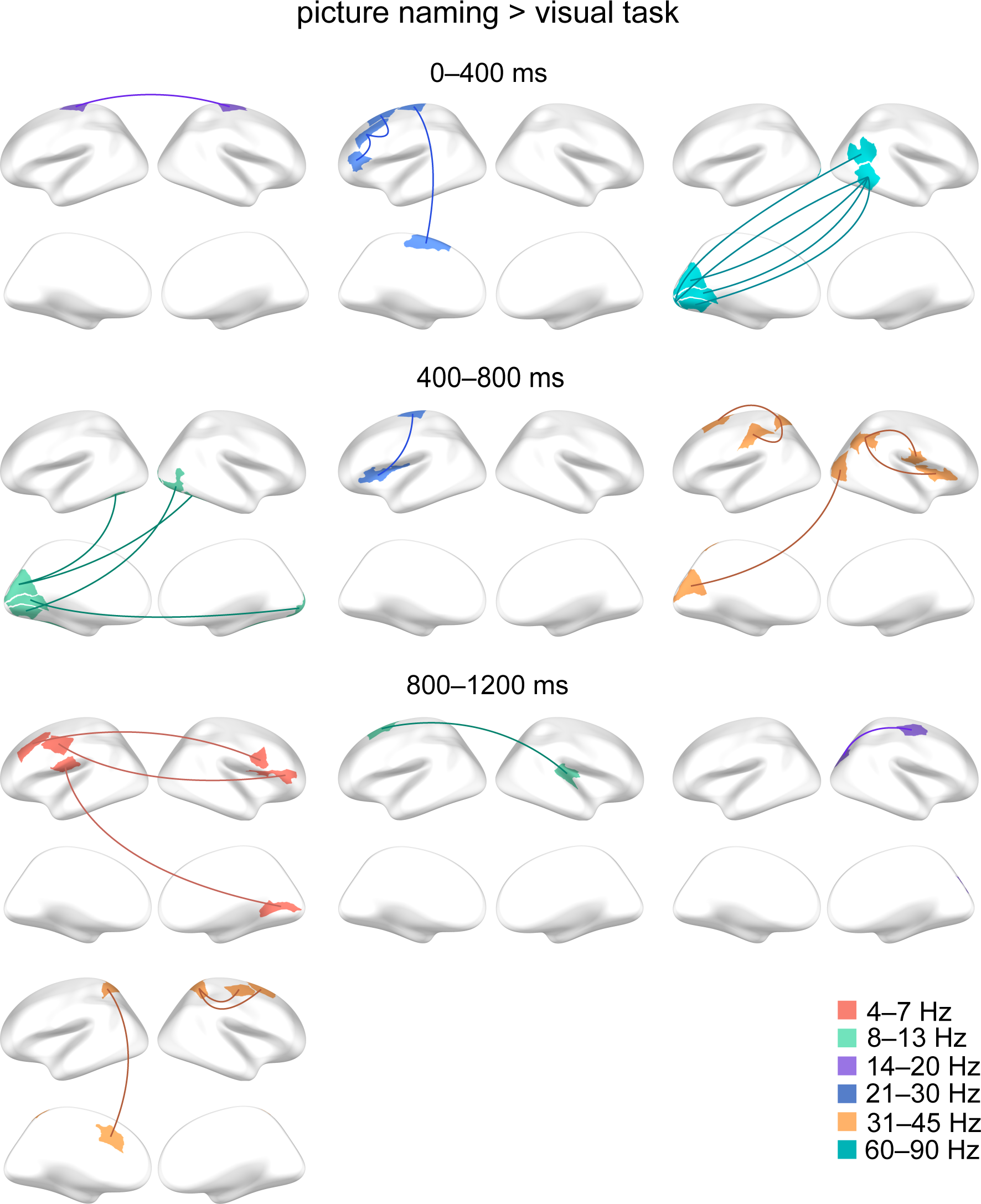
The consistent connections showing increased connectivity in the picture naming vs. the visual task. The consistent (ICC>0.4) connections are summarized on schematic brains (upper row lateral and lower row medial views) for different time windows and frequency bands. The statistical threshold in connection selection was p<0.0001.

Figure 4 illustrates decreased connectivity in the picture naming task compared to the visual task. A consistent naming-related connectivity decrease was observed most notably in the gamma band between the bilateral motor areas in the late time window (800–1200 ms). Consistent right-lateralized naming-related connectivity decreases were observed in multiple frequency bands and time windows: For instance, alpha connectivity decreased between right parietal and right temporal and parietal regions (400–800 ms). Beta connectivity decreased mainly between right frontal and right motor areas (400–800 ms) and between right parietal and right temporal regions (800–1200 ms).

**Figure 4.**
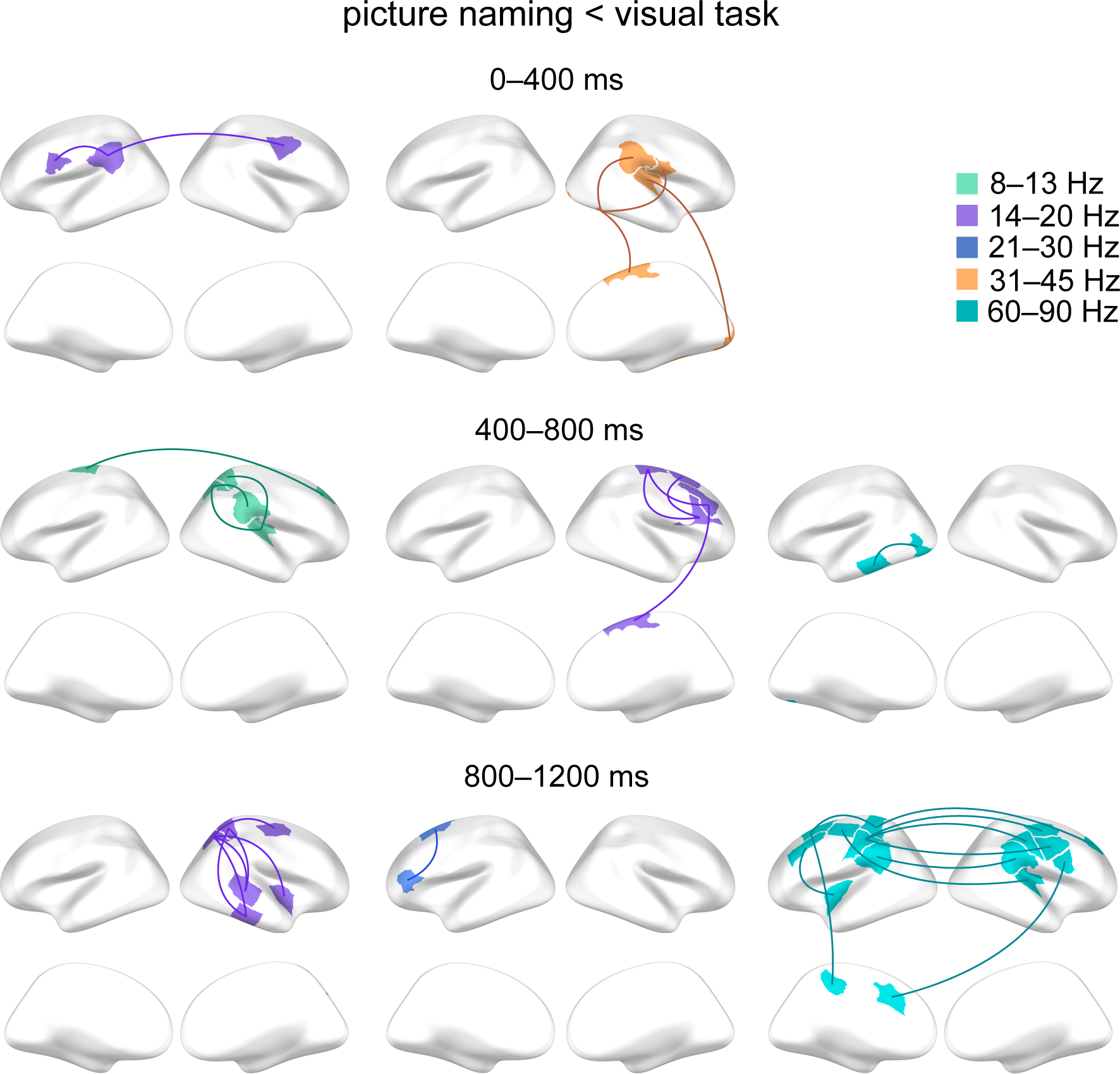
The consistent connections showing decreased connectivity in the picture naming vs. the visual task. The consistent (ICC>0.4) connections are summarized on schematic brains (upper row lateral and lower row medial views) for different time windows and frequency bands. Statistical threshold in connection selection was p<0.0001.

To summarize the laterality of the naming-related consistent connections, connectivity increases were detected in 14 bilateral, 8 left-lateralized, and 5 right-lateralized connections (Figure 5). Connectivity decreases were detected in 17 right-lateralized, 9 bilateral and 6 left-lateralized connections (Figure 5).

**Figure 5.**
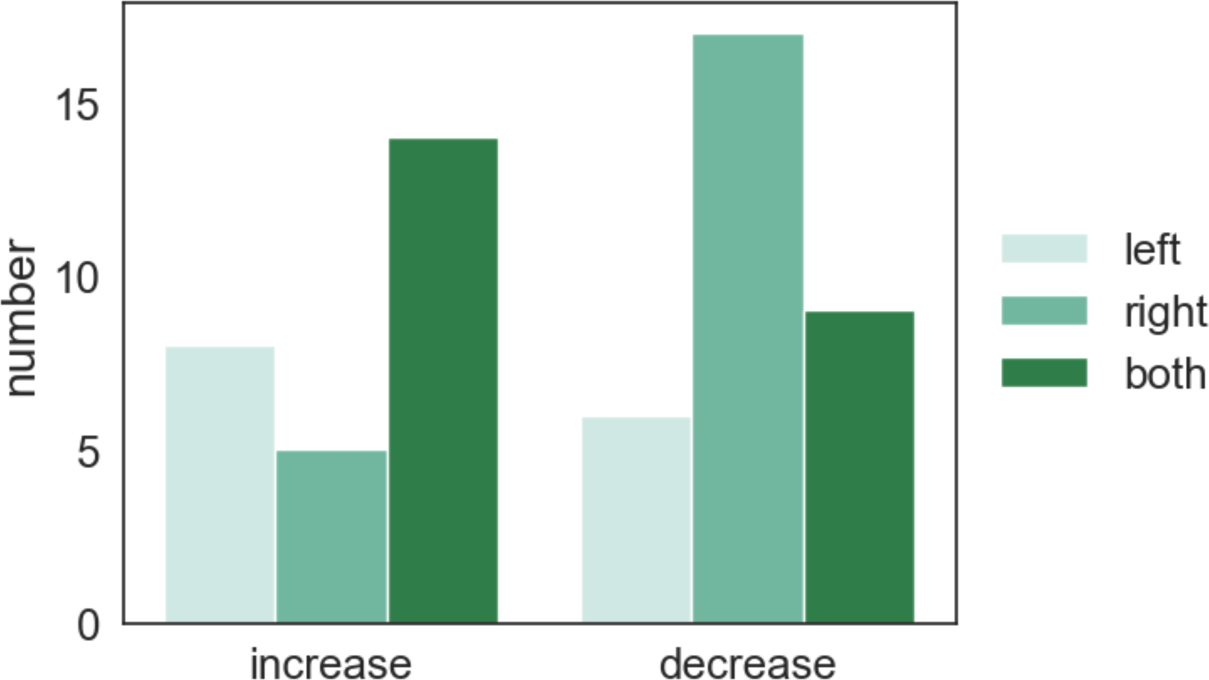
Connectivity modulation and laterality. Numbers of consistent naming-related connections grouped by connectivity increase and decrease and whether they were left-lateralized, right-lateralized, or bilateral (both).

### 3.3 Behavioral performance and consistency of connections

The consistent naming-related connections tended to be more frequently associated with language performance (Figures 6A and 6B) than the inconsistent connections (Statistical threshold in connection selection p<0.001: consistent 8.5 % and inconsistent 4.9 % connections, Chi-square test, p=0.0012. Statistical threshold in connection selection p<0.0001: consistent 10.2 % and inconsistent 4.6 % connections, Chi-square test, p=0.14.). In particular, consistent frontal connections were associated with language performance (Figure 6 C).

**Figure 6.**
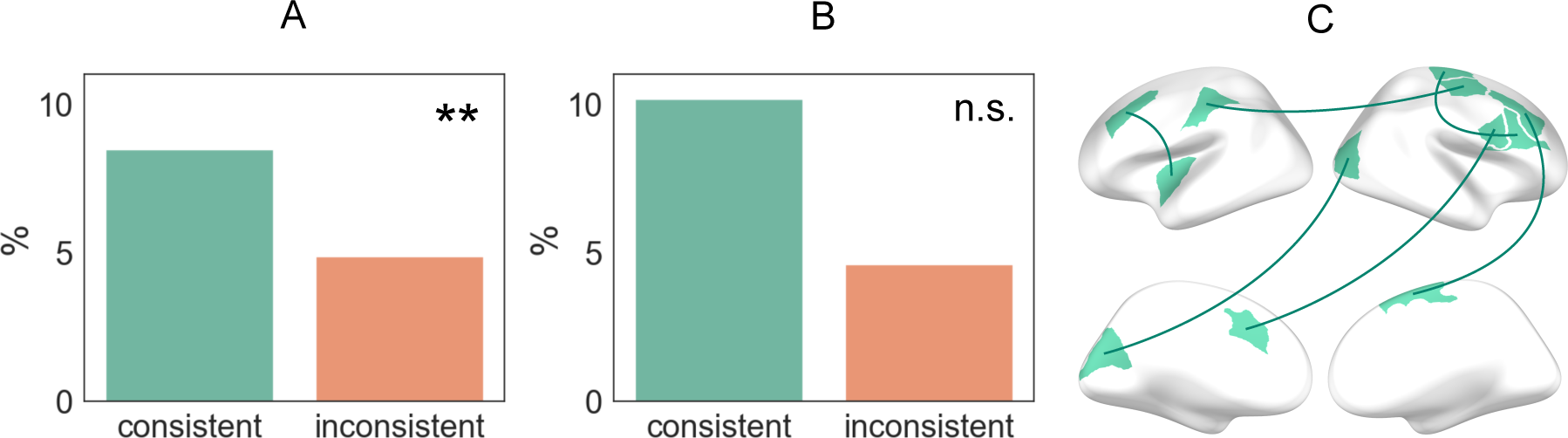
Language performance and consistency. The percentages of the connections for which the language performance explained (multiple regression model) connectivity strength for the consistent and inconsistent connections. Chi-square test: n.s.=non-significant, **p<0.01. Statistical threshold in connection selection p<0.001 (A) and p<0.0001 (B). C: The consistent connections that were associated with language performance (statistical threshold in connection selection p<0.0001).

### 3.4 MEG strength and structural connectivity related to consistency

The functional and structural properties underlying the consistent and inconsistent connections are illustrated in Figure 7. The consistent connections were functionally stronger (Figure 7: MEG strength) than the inconsistent connections (Statistical threshold in connection selection p<0.001: median consistent 3.4 and median inconsistent 2.9, Mann-Whitney U-test, p=0.0022. Statistical threshold in connection selection p<0.0001: median consistent 5.4 and median inconsistent 4.3, Mann-Whitney U-test, p=0.016.). The consistent connections were structurally stronger (Figure 7: number of direct streamlines) than the inconsistent connections (Statistical threshold in connection selection p<0.001: median consistent 186 and median inconsistent 84, Mann-Whitney U-test, p=1.2e-08. Statistical threshold in connection selection p<0.0001: median consistent 425 and median inconsistent 130, Mann-Whitney U-test, p=0.023.). The consistent connections were associated with shorter structural path lengths (Figure 7: shortest path length) than the inconsistent connections (Statistical threshold in connection selection p<0.001: median consistent 2.2e-04 and median inconsistent 3.9e-04, Mann-Whitney U-test, p=4.3e-07. Statistical threshold in connection selection p<0.0001: median consistent 1.7e-04 and median inconsistent 3.6e-04, Mann-Whitney U-test, p=0.0089.). The medians of the connections’ structural betweenness centrality were zero both for the consistent and the inconsistent connections (Figure 7: betweenness centrality).

**Figure 7.**
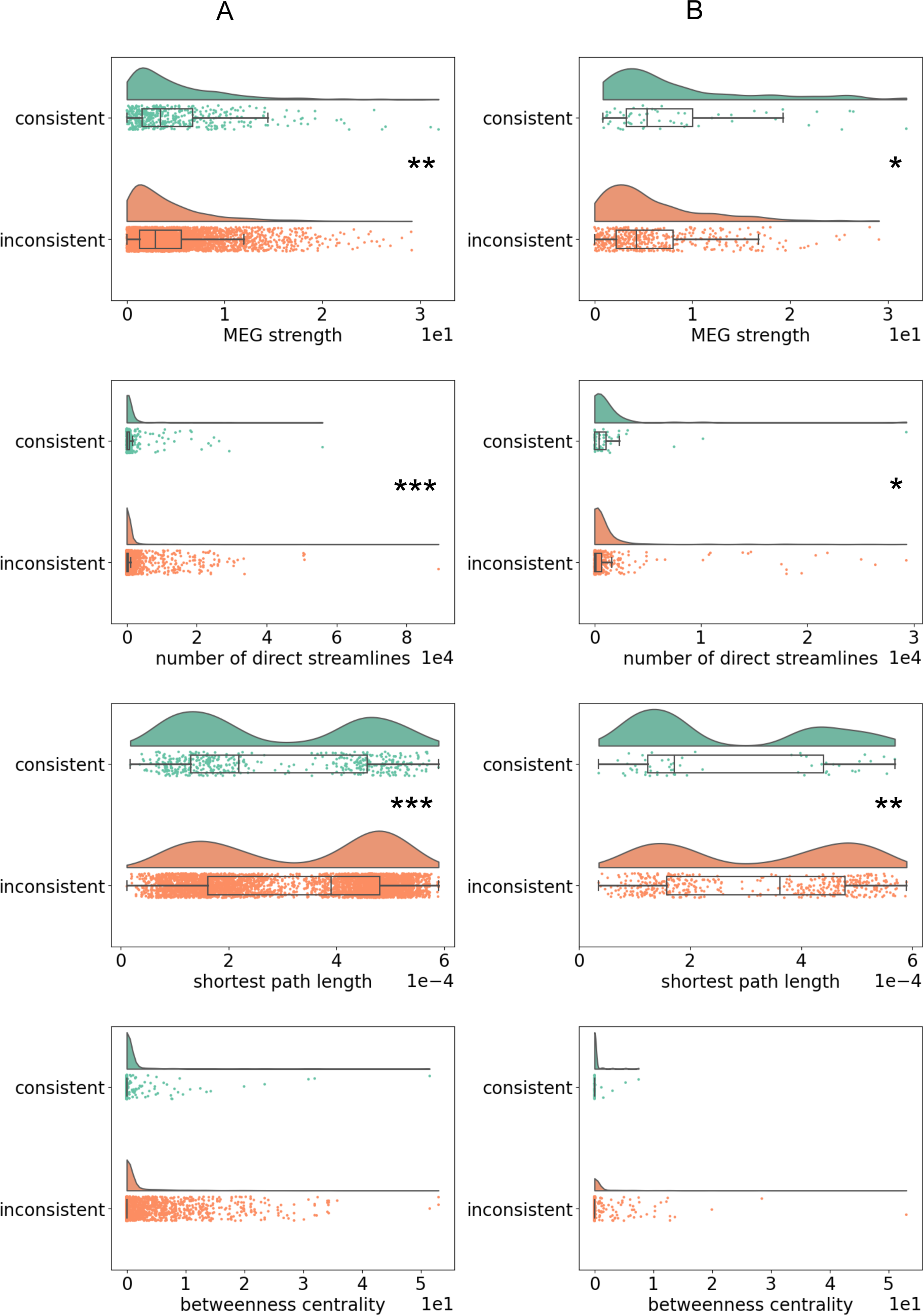
Functional and structural properties of the connections with respect to consistency of connectivity. The distributions, medians, and confidence intervals of the absolute MEG strength, number of direct structural streamlines, structural shortest path length, and structural betweenness centrality for the consistent and the inconsistent connections. Mann-Whitney U-test: *p<0.5, **p<0.01, ***p<0.001. Statistical threshold in connection selection p<0.001 (A) and p<0.0001 (B).

## 4 DISCUSSION

The present MEG study aimed to determine reliable dynamic functional connections related to picture naming and define behavioral, functional, and structural properties that may play a role in reliable functional connections. Altogether, we observed that connectivity related to picture naming was modulated compared to a visual task consistently and dynamically in different time windows, frequency bands and hemispheres as has also been seen in earlier connectivity studies related to language (Liljeström et al., 2015; Liljeström et al., 2018; Schoffelen et al., 2017). The connections that showed consistent task-related modulation across two days formed a fairly small subset of those connections that were selected for the consistency analysis. A particularly salient consistent pattern comprising naming-related beta connectivity increase in the left motor and frontal regions (0–400 ms and 400–800 ms) and gamma connectivity decrease in the bilateral motor regions (800–1200 ms) was detected which we interpret to represent the motor preparation of speech production. Interestingly, consistent connectivity decreases were emphasized in the right hemisphere. Moreover, we found that the consistent connections were more frequently associated with language tasks, were functionally and structurally stronger than the inconsistent connections, and were linked with structurally shorter path length.

### 4.1 Reliable connectivity related to picture naming

By focusing on time windows preceding overt speech output in a picture naming task, we here aimed to capture consistent connectivity modulations engaging brain processes related to preparation of speech production. In accordance with this aim, we detected a consistent naming-related connectivity pattern comprising beta connectivity increase in the left motor and frontal regions (0–400 ms and 400–800 ms) which we suggest represents motor preparation for speech production. Previously, connectivity modulation between left motor, premotor and frontal (Li et al., 2020) regions has been shown to play a role in language production, especially within the beta frequency band (Liljeström et al., 2015). We also detected stronger gamma connectivity during the visual task than during the naming task in the bilateral motor regions (800–1200 ms). Bilateral motor gamma connectivity during conditions when speech should be suppressed have been detected earlier and suggested to be involved in inhibition of speech production (Liljeström et al., 2015). Here, the observed consistent gamma connectivity pattern might represent the need to inhibit speech production during the visual task.

Several studies have shown that the right hemisphere plays a significant role during language tasks (Bozic, Fonteneau, Su and Marslen-Wilson, 2015; Chai et al., 2016; Doron, Bassett and Gazzaniga, 2012; Fedorenko, Hsieh, Nieto-Castañón, Whitfield-Gabrieli and Kanwisher, 2010; Medaglia et al., 2015). In the present study, we detected consistent right-lateralized alpha connectivity decrease (picture naming vs. visual task) between parietal and temporal and parietal regions (400–800 ms) and beta connectivity decrease between parietal and temporal regions (800–1200 ms). Similarly, earlier language studies have reported right-lateralized connectivity modulations between parietal and temporal regions. These modulations have been proposed to be involved in broader information of objects beyond language processing (Chai et al., 2016; DeSalvo et al., 2014). We suggest that the observed consistent connectivity decreases in the right parietal and temporal regions represent the processing of the visual image object during the baseline visual task. To summarize, the results of the present study showed that the consistent connectivity decreases were emphasized in the right hemisphere which might be due to the nature of the baseline visual task. This finding underlines the effect of the selected (baseline) task to the interpretation of the connectivity results.

### 4.2 Behavioral performance and reliability

When determining task-related connectivity, the validity of the results needs to be verified. One approach to validate the functional connectivity patterns is to relate them to behavior, since connections linked to behavior are likely to represent behaviorally meaningful and behavior-specific connections (Cohen, 2018; Jiang et al., 2020). Indeed, there are results showing that the modulation of functional connectivity is associated with behavioral performance (Bassett and Mattar, 2017; Elton and Gao, 2015; Finn et al., 2015; Jiang et al., 2020). For language processing, recent studies have emphasized the relevance of frontal connections in language performance (Mollo et al., 2016; Tomasi and Volkow, 2020; Vatansever et al., 2017; Yu et al., 2018). Similarly, we detected consistent frontal connections that were associated with language performance. We argue that this association provides additional validation for the current reliable language-related connectivity results.

There is a lack of research systematically studying the link between the behavioral utility of the connection and its consistency. One earlier study (Noble et al., 2017) reported that a connection’s consistency is not meaningfully correlated with the connection’s contribution to behavioral prediction, suggesting that the most reliable connections are not the most informative connections for behavior. In contrast, we observed that there was a tendency for consistent connections to be more frequently associated with behavioral performance. This discrepancy of findings remains to be further investigated in future studies with dedicated experimental designs. Indeed, one may ask whether it is meaningful to define task-related reliable connections if those would not have behavioral utility.

### 4.3 Functional and structural properties of the connections and reliability

Currently, little is known of what are the functional and structural properties of the reliable functional connections. Earlier findings indicate that stronger functional connections are more reliable (Garcés et al., 2016; Noble et al., 2019). Similarly, we observed that the consistent connections were functionally stronger than the inconsistent connections. However, even fewer studies have investigated what are the structural properties of the reliable functional connections.

Earlier studies have demonstrated similarities in the configuration of structural and functional connectivity, yet the relationship between the two is unknown (see for a review Avena-Koenigsberger et al., 2018). Structural connectivity is thought to constrain (Avena-Koenigsberger et al., 2018) and shape (Honey, Thivierge and Sporns, 2010) functional connectivity. For example, the strength of a functional connection between two brain areas depends on the strength of the structural connection between those areas (Hermundstad et al., 2013; Honey et al., 2009). Consequently, structural properties underlying functional connectivity may offer insight about neural communication between brain regions (Avena-Koenigsberger et al., 2018).

The present study investigated whether the number of direct structural streamlines and the structural length of the connection had an impact on the functional consistency. Currently, there is some evidence that structurally connected brain regions have more consistent functional connections than structurally unconnected brain regions (Griffa et al., 2017; Shen et al., 2015). The results of the present study further demonstrated that the number of direct streamlines were greater for consistent than inconsistent connections. Our results also showed that the consistent connections were associated with shorter structural path length than the inconsistent connections. The shortest path length is the topological distance between two brain regions. Minimizing the path length is desirable since longer paths are more susceptible to, e.g., noise and longer transmission delays (Avena-Koenigsberger et al., 2018). For these reasons, shortest paths are thought to ensure reliable and efficient communication between brain regions (Avena-Koenigsberger et al., 2018). We suggest that our findings indicate that the most efficient communication between brain regions is emphasized in consistent connections.

### 4.5. Methodological considerations and future directions

#### 4.5.1 Coherence as a connectivity measure

In this study coherence was used as a connectivity measure. Coherence emphasizes the signal transmission along paths that can be dynamically reconfigured and is, thus, an excellent connectivity measure to study task-related modulations of connectivity (Fries, 2005; Liljeström et al., 2015; Saarinen, Jalava, Kujala, Stevenson and Salmelin, 2015). However, coherence is prone to field spread effects, which has to be taken into account. There are some other connectivity measures that are less sensitive to field spread, such as imaginary coherence. Nevertheless, to use imaginary coherence to study task-related modulations of connectivity can be problematic (Gross et al., 2013). To summarize, the connectivity results are dependent on the chosen connectivity measure and every measure has advantages and disadvantages which have to be evaluated and addressed. Here we chose coherence as a connectivity metric, since it is particularly well-suited to study dynamic task-related modulations of connectivity. The best effort was made to minimize the effects of field spread (see sections 2.4 MEG preprocessing and connectivity analysis and 2.6 Power difference between the tasks).

#### 4.5.2 The selection of connections for the consistency analysis

When determining the consistent connections, the selection of the connections that are being studied has its effect on the results. In this study, we did not use any priors when selecting the studied connections, rather the selection was based on the connectivity data itself. We used two statistical thresholds when selecting the connections for the consistency analysis. This selection had an impact on the number of the consistent connections. However, the selection did not affect the general outcomes of how the consistent connections were associated with the behavioral performance and functional and structural properties of the connections.

#### 4.5.3 How to improve consistency?

The results of this study illustrated that the percentage of the consistent connections of all the selected connections is quite modest. This finding is in line with an earlier meta-analysis of 25 fMRI studies showing that, on average, individual connections exhibit low reliability (ICC 0.29) (Noble et al., 2019). This raises a question: What should be done in future connectivity studies to improve reliability? The results of this study provide some hints that it might be beneficial to utilize behavior-based validation as well as functional and structural information when defining reliable connections.

### 4.6. Conclusions

To gain a more accurate and comprehensive view of the dynamic nature of brain networks, we need to be able to define reliable and task-related brain networks. Overall, this study defined reliable dynamic functional connectivity related to language and also added to the understanding of what makes a connection reliable.

## ACKNOWLEDGMENTS

We acknowledge the computational resources provided by the Aalto Science-IT project. We thank Susanna Aro for helping to implement the functional connectivity codes for the data set.

## SUPPORTING INFORMATION

The raw or processed MEG or MRI data from individual subjects cannot be made openly available, according to the ethical permission and national privacy regulations at the time of the data collection. The derived results that served as input for the figures will be made openly available.

Analysis methods used in the study are based on openly available software packages for MEG and MRI analysis, as described in the Materials and Methods section. The DICS beamformer as described in van Vliet et al. (2018) has been incorporated into MNE Python (https://mne.tools) version 0.16 and above, and the methods that were used for analysis of connectivity are available in the ConPy package (https://github.com/AaltoImagingLanguage/conpy). The graph theoretical metrics from diffusion MRI data were calculated with the Brain Connectivity Toolbox (Rubinov and Sporns, 2010).

